# Effector T Cells under Hypoxia have an Altered Transcriptome Similar to Tumor-Stressed T Cells Found in Non-Responsive Melanoma Patients

**DOI:** 10.1101/2024.09.27.615474

**Authors:** Mate Z. Nagy, Lourdes B. Plaza Rojas, Justin C Boucher, Elena Kostenko, Anna L. Austin, Ahmad Tarhini, Zhihua Chen, Dongliang Du, Awino Maureiq E. Ojwang, Joshua Davis, Katarzyna A. Rejniak, Timothy I. Shaw, Jose Alejandro Guevara

## Abstract

**Background:** In the Tumor Microenvironment (TME), hypoxia stands as a significant factor that modulates immune responses, especially those driven by T cells. As T cell-based therapies often fail to work in solid tumors, this study aims to investigate the effects of hypoxia on T cell topo-distribution in the TME, gene expression association with T cell states, and clinical responses in melanoma.

**Methods:** To generate detailed information on tumor oxygenation and T cell accessibility, we utilized mathematical modeling of human melanoma tissue microarrays (TMAs) that incorporate oxygen supply from vessels, intratumoral diffusion, and cellular uptake. We created tumor maps and derived plots showing the fraction of CD4 and CD8 T cells against the distance to the nearest vessel and oxygen pressure. To assess their function and transcriptional changes caused by hypoxia, effector T cells were generated and cultured under hypoxia (0.5% oxygen) or normoxia (21% oxygen). The T cell hypoxia-transcriptional signature was compared against datasets from msigDB, iATLAS (clinical trials of melanoma patients treated with immune checkpoint inhibitors), ORIEN AVATAR (real-world melanoma patients treated with immune checkpoint inhibitors, ICIs), and a single-cell atlas of tumor-infiltrating lymphocytes (TILs).

**Results:** We made three specific observations: 1) in melanoma T cells preferentially accumulated in oxygenated areas close to blood vessels (50-100 micrometers from the vasculature in the regions of high oxygen availability) but not in hypoxic areas far from blood vessels. 2) Our analysis confirmed that under hypoxia, T cell functions were significantly reduced compared to normoxic conditions and accompanied by a unique gene signature. Furthermore, this hypoxic gene signature was prevalent in resting and non-activated T cells. Notably and clinically relevant, the hypoxic T cell gene set was found to correlate with reduced Overall Survival (OS) and reduced progression-free survival (PFS) in melanoma patients, which was more pronounced in non-responder patients undergoing ICI therapy. 3) Finally, compared with a single-cell atlas of tumor-infiltrating T cells, our hypoxia signature aligned with a population of cells at a state termed stress response state (T_STR_).

**Conclusion:** Our study highlights the critical role of hypoxia in shaping T cell distribution and its correlation with clinical outcomes in melanoma. We revealed a preferential accumulation of T cells in oxygenated areas. Moreover, hypoxic T cells develop a distinct hypoxic gene signature prevalent in resting, non-activated T cells and T_STR_ that was also associated with poorer outcomes, particularly pronounced among non-responders to ICIs.

**Key Messages:** **What is already known on this topic:** Hypoxia significantly impairs T cell functions, including reduced proliferation and cytokine production. This impairment may contribute to immune evasion and resistance to immune therapies, such as ICIs, adoptive transfer of TILs, and Chimeric Antigen Receptor (CAR) T cells. Despite the established impact of hypoxia on T cell function, the precise spatial distribution of T cells in relation to oxygen availability within the TME and how this affects clinical outcomes in melanoma patients remains unclear. Additionally, the specific transcriptional changes in T cells induced by hypoxia and their prevalence in different T cell states, as well as the implications for resistance to ICIs, have not been thoroughly investigated. Understanding these aspects is crucial for developing targeted therapies to overcome hypoxia-induced resistance and improve immunotherapy efficacy.

**What this study adds:** This study elucidates the profound impact of hypoxia on T cells in melanoma. Our findings reveal that T cells accumulate in well-oxygenated regions near blood vessels, whereas hypoxic conditions significantly impair their proliferation and cytokine production. By addressing the transcriptional changes induced by hypoxia, we demonstrated its prevalence in resting and non-activated T cells and T_STR_. Moreover, we found that hypoxia is associated with shorter OS and PFS in melanoma patients, particularly in non-responders to ICIs. This research highlights the critical role of hypoxia in modulating T cell spatial behavior, contributing to immune evasion and poor clinical outcomes in melanoma.

**How this study might affect research, practice, or policy:** The findings emphasizes hypoxia’s critical role in modulating T cell behavior and its potential as a biomarker for treatment outcomes. These insights could inform the development of therapeutic strategies aimed at improving T cell function in hypoxic TMEs, thereby enhancing the efficacy of immunotherapies for melanoma and other solid tumors. Additionally, the study’s results may influence policies regarding the evaluation and implementation of combination therapies that target hypoxia to improve patient outcomes.

## Introduction

Chaotic blood vessel formations and drastic differences in oxygen availability are hallmarks of solid tumors.^1^ Oxygen concentrations in these tumors can range from 0.3 to 4 mmHg O2, significantly lower than the 20-100 mmHg O2 found in normal tissues, resulting in a hypoxia environment.^2^ Under these low-oxygen conditions, cancer cells express pro-survival genes, enhancing angiogenesis and metabolically adapting to oxygen deprivation.^3^ This adaptation leads to increased cellular proliferation, invasion, treatment resistance, and worsening prognosis.^4^ In contrast, hypoxia exerts a suppressive effect on T cell properties.^5–9^ These studies emphasize the need to better understand hypoxic TMEs and find reliable tools to identify T cell populations that are better suited to target solid tumors.

Studies have demonstrated relationship between the type, density, and location of TILs and clinical outcomes in colorectal cancer patients.^10^ Moreover, gene expression profiling has emerged as a significant tool in melanoma research, enabling the identification of specific patterns linked with melanoma progression and prognosis.^11, 12^ The tumor mutation burden is another promising biomarker, often associated with a better response to immunotherapy.^13^

While cancer cells thrive under hypoxic conditions, T cells become highly dysfunctional.^14^ Hypoxia suppresses T cell cytotoxicity and promotes regulatory T cell recruitment, further dampening the anti-tumor immune response.^15^ This dual effect of hypoxia—enhancing cancer cell survival while impairing T cell function—presents a significant challenge in developing effective cancer immunotherapies.

We hypothesized that T cell distribution in the tumor would depend on oxygen accessibility. Our data indicates that in melanoma, oxygen accessibility determines T cell topo-distribution within the TME, function and transcriptional program. We focused on highlighting the effects of hypoxia exposure on T cells, both functionally and transcriptionally, and how these associate with clinical responses. Using RNA sequencing of effector T cells, we uncovered a hypoxia-gene signature predominantly present in inactive T cells. This signature correlates with decreased survival in melanoma patients and in those unresponsive to ICI therapy. We compared our gene signature with an extensive single-cell atlas of T cells and found that our hypoxia-associated signature is enriched in tumor-stressed T cells.^16^

Our findings further emphasize the significant role of oxygen accessibility in modulating T cell distribution, function, and behavior within the TME and how these factors may affect clinical responses. By understanding the relationship between hypoxia and T cell activity, we can better identify which T cell populations are most effective in targeting tumors. This knowledge may pave the way for developing improved immune therapeutics that select for or engineer T cells capable of thriving in hypoxic conditions or that are not yet genetically altered by hypoxia. Our results highlight the potential for targeting hypoxia-associated pathways to enhance the efficacy of immunotherapies in treating solid malignancies.

## MATERIALS AND METHODS

### Immunohistochemical analysis and computational modeling of human melanoma tissue microarray (TMA)

The study utilized a human TMA containing 31 biopsy samples (cores and periphery) from 17 melanoma patients treated at Moffitt Cancer Center, representing different regions of melanoma tumors. The study was conducted under the approval of the IRB at Moffitt Cancer Center, ensuring adherence to ethical guidelines and patient consent protocols. TMA blocks were sectioned into 5 µm thick slices using a microtome and mounted onto Superfrost Plus microscope slides (Thermo Fisher Scientific). Slides were deparaffinized in xylene and rehydrated through a graded series of ethanol, followed by rinsing in distilled water. For antigen retrieval, sections were immersed in citrate buffer (pH 6.0), heated in a microwave oven, and cooled at room temperature. The sections were stained for lymphocytes (CD3, CD8, and CD4), cell nuclei (DAPI), and vasculature (CD31). The stained TMAs were scanned and saved as high-resolution images. These images were segmented using QuPath software^17^ to identify individual cell positions and areas. In addition, an in-house MATLAB^®^ (The MathWorks Inc.) routine was used to determine the locations of all capillaries. The spatial distribution of oxygen in the TMA tissue was modeled based on physics first principles, as a balance between oxygen supply from the vasculature, its intratumoral diffusion, and uptake by the cells located within the tissue, following our previous work.^18–20^ The stable oxygen distribution was achieved computationally by iterative calculation of the reaction-diffusion equation (eq.1) until the L_2_ norm between two consecutive oxygen distributions fell below the given threshold, i.e., 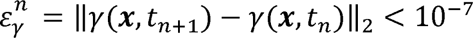, where the change in oxygen *γ*(***x***,*t*) was given by:

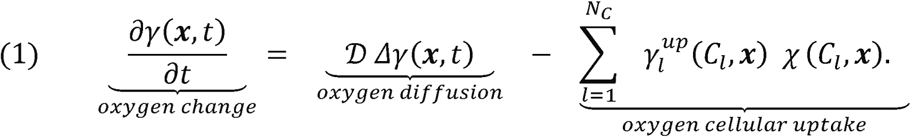

Here, the oxygen diffusion coefficient was *D* = 100*μm*^2^/*s*, and the cell neighborhood was defined by eq.(2):

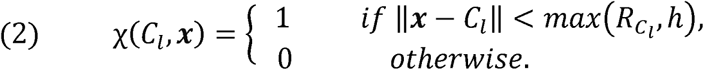

The oxygen uptake by the cell 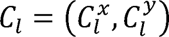 of radius *R_C_l__* was defined using the Michaelis-Menten kinetics (eq.3) with constants *V_m_* = 0.62143σ*g*/*s*, and *k_m_* = 1.344σ*g*/*μm*^3^ (scaling parameter σ*g* = 0.05*ag*), and the grid size *h* = 2*μm*,

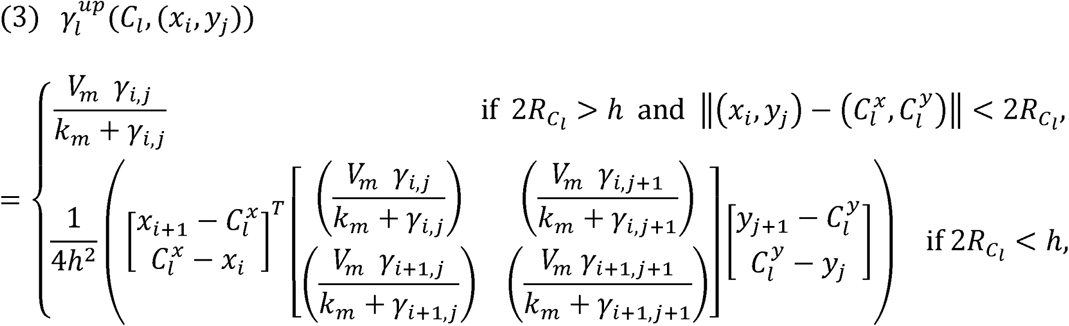

where 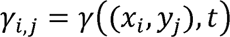. The oxygen supply at every vasculature location was *γ_max_* = 60 *mmHg*.

For the given TMA melanoma tissue with calculated stable oxygen distribution, two metrics were computed for each T cell: the distance to the nearest vasculature location and the local averaged oxygen level surrounding the cell. These two measurements were used to characterize the regions of melanoma tissue occupied by the T cells. These data were used to generate detailed tumor oxygenation and T cell accessibility maps, showing the oxygen levels experienced by each T cell and their distances from the closest vessel.

### Generation of effector T cells from healthy donors and assessing proliferative and functional capacities under hypoxia

Cryopreserved healthy human peripheral blood mononuclear cells (PBMCs) were obtained from AllCells. Cryopreserved samples were activated using anti-human CD3 OKT-3 clone from inVivoMAb (catalog: BE0001-2) at a concentration of 20 ng/mL, along with 300 IU/mL of teceleukin (recombinant interleukin-2) from Hoffmann-LA Roche. The cells were cultured at 37°C in a 5% CO2 incubator for 6 days using Corning RPMI 1640 supplemented with 5% fetal bovine serum (Sigma-Aldrich) and 5% penicillin-streptomycin (ThermoFisher). During culture, activated cells received 60 IU/mL of IL-2 as needed when the media was changed. After six days of culture, the cells were washed, counted, and stained following Invitrogen’s CellTraceTM Violet Stain protocol. A portion of the cells were used for baseline CellTrace Violet (CTV) measurements. The cells were also stained with Invitrogen’s near-IR fluorescent reactive dye and an anti-human CD3 dye (UCHT1 clone) from BioLegend. Stained cells were analyzed using a FACSymphony benchtop analyzer. The remaining cells were split equally and cultured under two conditions: 37°C, 5% CO2, and 21% oxygen (normoxia) or 37°C, 5% CO2, and 0.5% oxygen (hypoxia). We utilized a BioSpherix C-Chamber system. This hypoxia chamber allows precise control of oxygen levels to simulate the TM). Cell cultures were maintained in complete RPMI, and flow cytometry analysis was performed at 48, 96, 168, and 216 hours after initial CTV staining. Supernatants from cell cultures on days 8, 11, 13, and 15 were collected for IFN-γ and TNF-α ELISA assays using MAPTECH ELISA Flex kits and protocols, analyzed with a Bio-Rad xMARKTM Microplate Spectrophotometer. Flow cytometry data was analyzed using FlowJo software version 10.8.2. Graphs were generated using GraphPad Prism software version 9.5.1 (528)

### RNAseq, pathway enrichment, and survival analysis

Read adapters were detected using BBMerge (v37.02)^21^ and subsequently removed with cutadapt (v1.8.1).^22^ Processed raw reads were then aligned to the human genome GRCH38 using STAR (v2.7.7a).^23^ Gene expression was evaluated as read count at the gene level with RSEM (v1.3.0) [PMID: 21816040] and the Gencode gene model v30. Gene expression data were normalized, and differential expression analysis between hypoxia and normoxia conditions was performed using DESeq2.^24^ A gene signature of upregulated hypoxia genes was defined based on a log2 fold change cutoff of 2 and an adjusted p-value cutoff of 0.05. Pathway enrichment was analyzed with GSEA based on signal-to-noise.^25^ Single-sample GSEA was calculated using the DRPPM-EASY app.^26^ Survival analysis was performed by PATH-SURVEYOR.^27^ The pan-cancer T cell atlas of single-cell RNAseq data, including UMAP coordinates, cell type annotations, and transcript expression values, were downloaded from https://singlecell.mdanderson.org/TCM/ and visualized by ISCVA.^28^ Hypoxia scores were calculated as the mean expression of genes designated as upregulated in the “hypoxia 0.1% T cell” signature.

## RESULTS

### Spatial Distribution of T Cells in Human Melanoma Correlates with Oxygen Accessibility and Proximity to Blood Vessels

Given that hypoxia is known to impair T cell functions,^6–9^ and considering that the spatial distribution of T cells in tumors and their relationship to oxygen availability are not well understood, we tested the hypothesis that T cell distribution within the tumor is influenced by oxygen accessibility. We stained 31 TMAs from 17 patients for CD31, CD4, CD8, and CD3, as illustrated in Figure 1A. Upon initial observation, it is apparent that the distribution of T cells within the TME is not homogeneous but is segregated to specific areas. To understand this non-random distribution, we focused on the T cells’ proximity to blood vessels. We examined cells expressing CD31, also known as Platelet Endothelial Cell Adhesion Molecule-1 (PECAM-1), a protein found on the surface of endothelial cells lining the interior surface of blood vessels. This allowed to determine proximity to the vasculature. An example of the individual cell marker staining are shown (Figure 1A).

**Figure 1:**
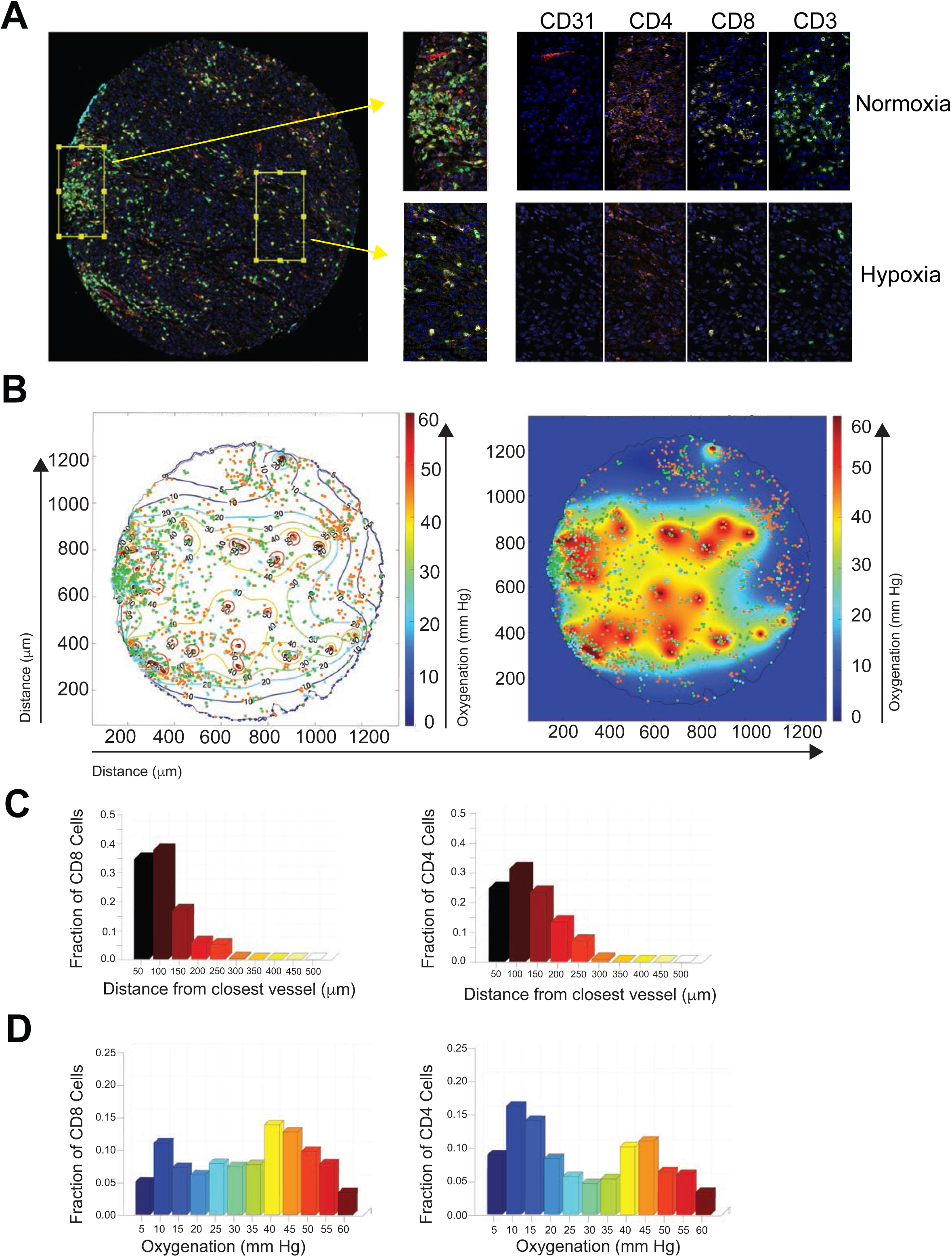
Spatial Distribution of T Cells in Human Melanoma Correlates with Oxygen Accessibility and Proximity to Blood Vessels. **(A)** Representative images of human melanoma TMAs stained with CD31 (vasculature), CD4, CD8, and CD3 (T cells). **(B)** Corresponding computationally-reconstructed oxygenation map of TMA with CD3 in light blue dots, CD8 T cells shown as green dots, and CD4 T cells as orange dots. Within the TME, the dark blue regions indicate hypoxic conditions (≤5 mmHg), while orange and red regions indicate well-oxygenated conditions (≥40 mmHg). **(C)** Quantification of the fractions of CD4 and CD8 T cells relative to their distance from the nearest blood vessel and **(D)** corresponding oxygen pressure sensed by the CD8 and CD4 cells.

Using coordinates of individual cells and vessels in each TMA, we generated tumor oxygenation and T cell accessibility maps (Figure 1B) as a balance between oxygen supply from the vessels, intratumoral diffusion, and cellular uptake. Next, we determined the level of oxygen that each T cell is experiencing, as well as the distances between T cells and their closest vessel, plotted as the fraction of CD4 or CD8 T cells against the distance to the closest vessel and oxygen pressure (Figure 1C). Analysis of these results showed that that the majority of CD4 and CD8 T cells (72% and 55.6%, respectively), were accumulated within 50-100 micrometers from the vasculature, indicated by dark red in Figure 1C. Moreover, the fraction of T cells under hypoxic conditions (≤5 mmHg), shown in dark blue, is less than the fraction of T cells in well-oxygenated conditions (≥40 mmHg), shown in orange and red (Figure 1D), namely 5% vs. 33.8% for CD8 cells and 8.9% vs. 26.8% for CD4 cells. Next, we evaluated these metrics in the rest of the melanoma TMAs separated by tumor core and periphery. We observed that cumulatively in all TMAs, CD8 and CD4 T cells were localized close to the blood vessels (76.3% of CD8 and 68.9% of CD4 in the core TMAs and 62.3% of CD8 and 50.6% of CD4 in periphery TMAs), where there is maximum oxygen above 40 mmHg (Figure 2A, B). We found no apparent differences between the localization of T cells at the core (Figure 2A) and periphery (Figure 2B). The individual TMA analysis is show in Figure S1. Overall, these findings suggest that T cells in melanoma tissues are predominantly situated close to blood vessels, highlighting the importance of oxygen accessibility in T cell distribution within the tumor microenvironment.

**Figure 2:**
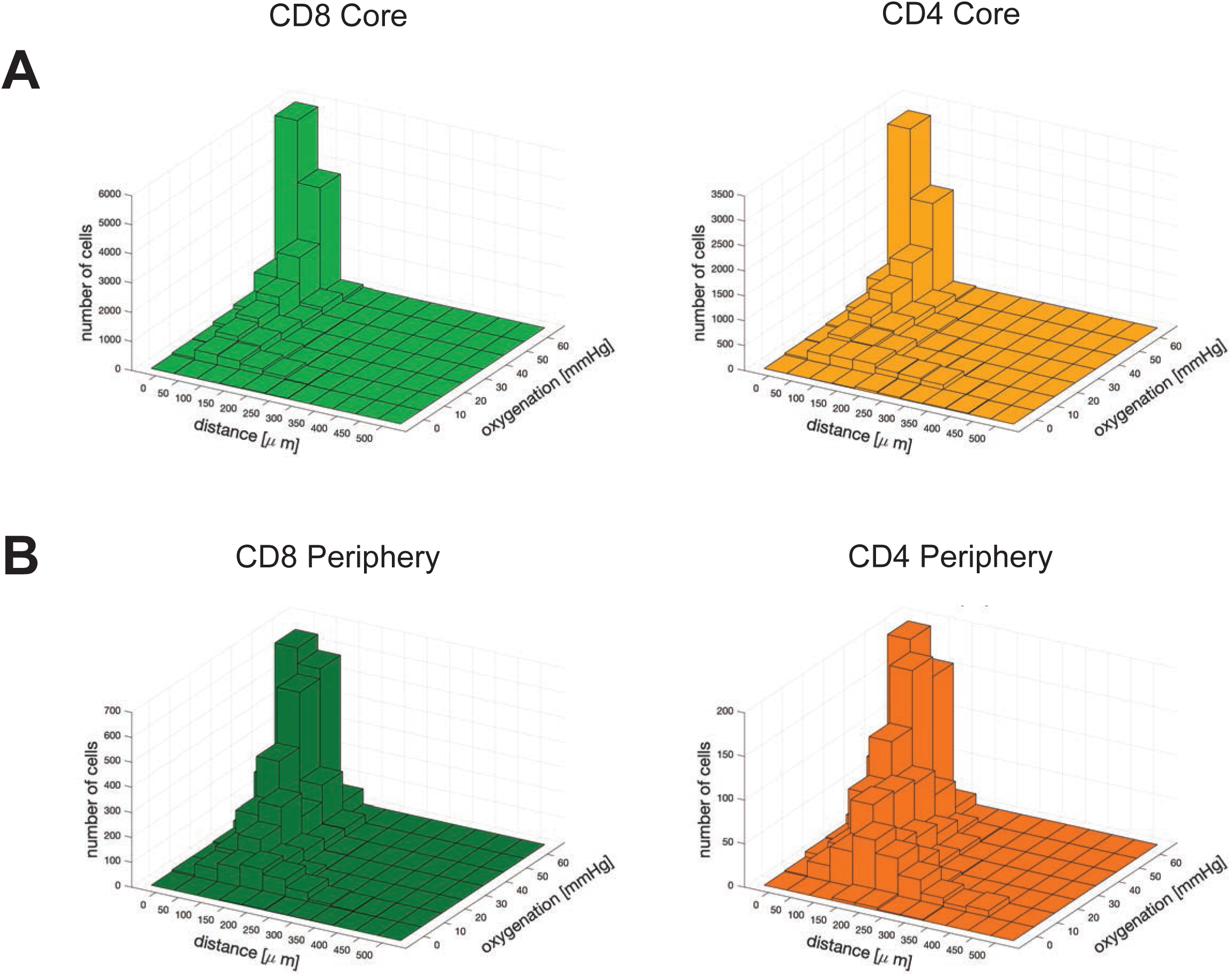
T Cell Localization and Oxygenation are Similar in TMAs of Tumor Cores and Peripheries. **(A)** Core: Quantification of CD8 and CD4 T cell distributions across all TMAs—core sections relative to the distance from the nearest blood vessel and corresponding oxygen pressure in tumor cores. (**B)** Peripheral: CD8 and CD4 T cell distributions in the TME—periphery relative to the distance from the nearest blood vessel and corresponding oxygen pressure in tumor peripheries.

### T cell proliferation and cytokine secretion is impaired under Hypoxic Conditions

The data above indicate that oxygen accessibility in the TME and T cell distribution are related. To further determine the impact of oxygen access, we activated T cells from healthy donors using anti-CD3 mAb and IL-2 for 6 days. The T cells were then exposed to a hypoxic condition with 0.5% oxygen or maintained in normoxic conditions with 21% oxygen for 3 days. We examined T cell proliferation using CTV tracer and found that an average of 23% of T cells proliferated under hypoxia compared to an average of 74% under normoxia (Figure 3A). Cell counting on days 6, 10, and 15 further validated these findings with normoxic T cells having a significant increase in numbers compared to hypoxic T cells at day 15 (Figure 3B). We also measured IFN-γ secretion after re-stimulation with plate-bound anti-CD3 mAb on days 8, 11, 13, and 15 and found that T cells under hypoxia produce significantly less INF-γ compared with T cell under normoxia, (Figure 3C). Taken together, these data indicate that hypoxia hinders the functional capacity of T cells, such as their ability to proliferate and produce cytokines.

**Figure 3:**
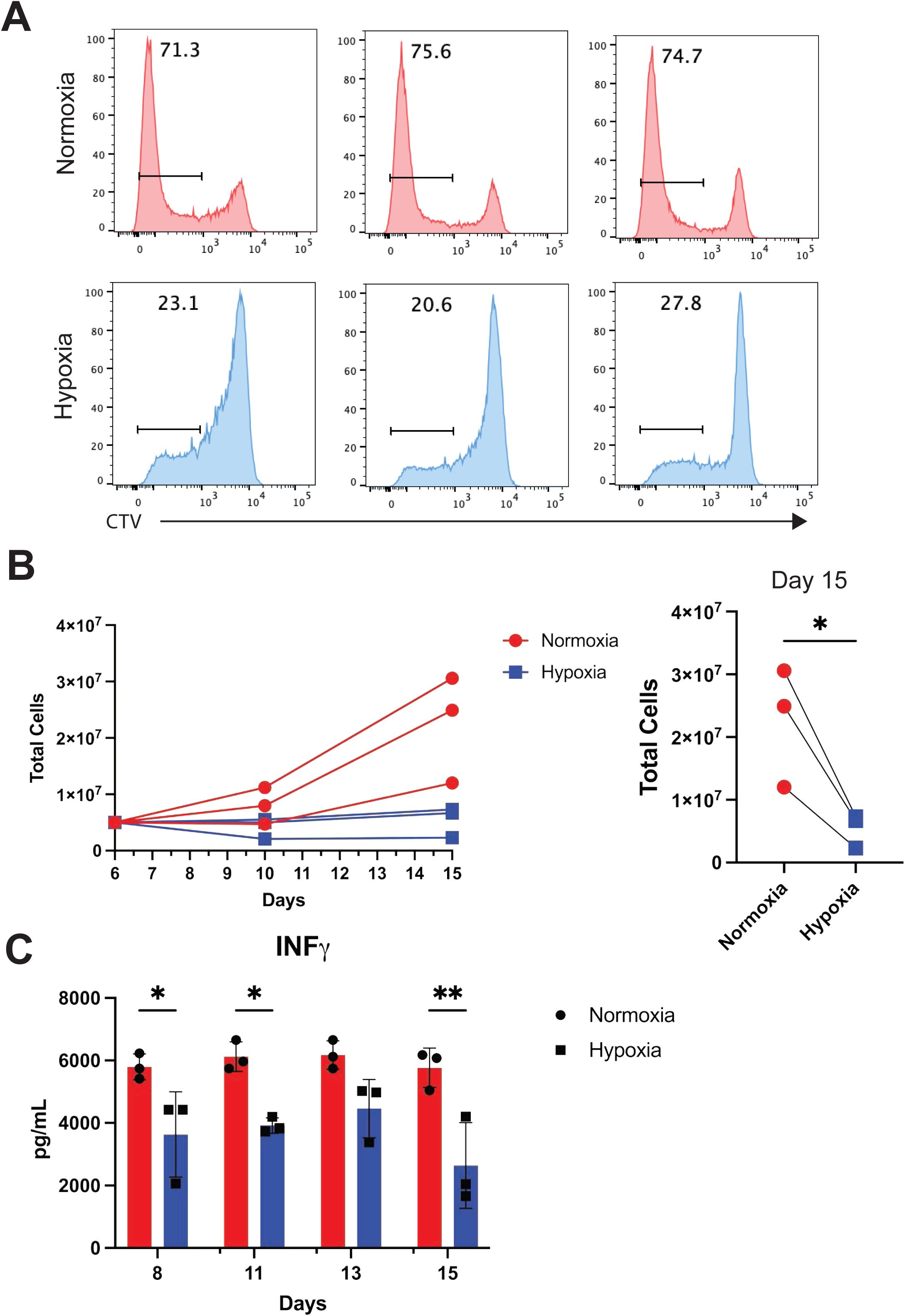
T cell proliferation and cytokine secretion are impaired under Hypoxic Conditions. **(A)** Cell proliferation of human T cells from three healthy donors under normoxia and hypoxia. T cells activated with anti-CD3 mAb and IL-2 for 6 days, then cultured under normoxic (21% O□) or hypoxic (0.5% O□) conditions for 3 days. Cell proliferation was assessed using CTV staining. **(B)** Cell counts on days 0, 10, and 15 under normoxia and hypoxia. **(C)** IFN-γ secretion on days 8, 11, 13, and 15 after re-stimulation with plate-bound anti-CD3 mAb.

### Hypoxic T cell gene expression is similar to resting T cells

Having demonstrated that oxygen availability dictates the functional capacity of T cells and associates with their spatial distribution within the TME, we sought to determine the impact of oxygen accessibility on the transcriptional program of T cells. To do this, we examined the transcriptional alterations in effector T cells, specifically focusing on how varying oxygen levels influence gene expression. We used effector T cells exposed to normoxia and hypoxia for 48 hour. We found differential gene expression patterns between T cells in hypoxic versus normoxic environments (Figure 4A). We identified a set of genes that were significantly reduced under hypoxia compared to normoxia forming a T cell hypoxia gene signature (Figure 4B). Next, when we compared these patterns against a human gene set that differentiates resting and activated CD8 and CD4 T cells.^29, 30^ We found that the gene expression profile for T cells under hypoxic conditions is enriched for genes typically upregulated in resting CD4 T cells (Figure 4C) and CD8 (Figure 4E). This enrichment starkly contrasts with the gene expression observed in normoxic samples (Figures 4D and 4F), suggesting that T cells in hypoxic conditions adopt a gene expression pattern akin to resting or quiescent T cells, as opposed to those in normoxic conditions. This demonstrates that the shift in the T cell signature under hypoxia substantially impacts their transcriptional activity. The data also shows that limited access to oxygen can significantly change T cell gene expression, which could have major implications for their clinical effects.

**Figure 4.**
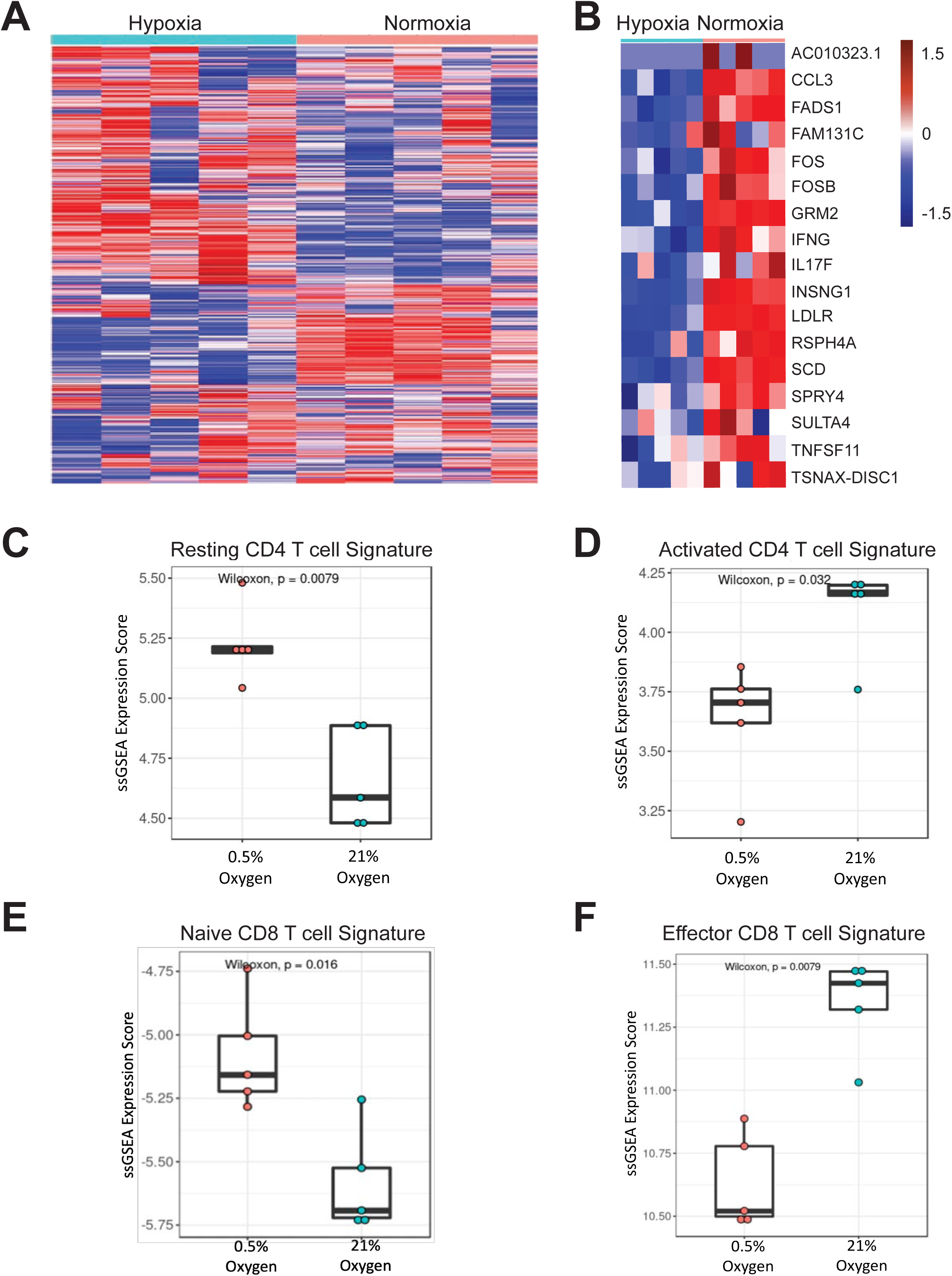
Hypoxia Induces a Resting-Like Gene Expression Profile in T Cells. **(A)** Gene expression patterns in effector T cells exposed to normoxic versus hypoxic conditions, illustrating the overall changes in gene expression profiles when T cells are subjected to hypoxic (0.5% oxygen) compared to normoxic (21% oxygen) environments. **(B)** Genes significantly reduced in effector T cells under hypoxia compared to normoxia. **(C)** T cell hypoxia gene signature enrichment in resting CD4 T cells. The gene expression profile for T cells under hypoxic conditions is enriched for genes typically upregulated in resting CD4 T cells, suggesting a shift towards a more quiescent state. **(D)** T cell hypoxia gene signature enrichment in activated CD4 T cells. Under normoxic conditions, there is a higher expression of genes associated with activated CD4 T cells, contrasting with the hypoxic condition which does not show this activation signature. **(E)** T cell hypoxia gene signature enrichment in naive CD8 T cells. Under hypoxic conditions, there is an enrichment of genes typically upregulated in naive CD8 T cells, again pointing towards a less active, more resting-like state. **(F)** T cell hypoxia gene signature enrichment in effector CD8 T cells. The gene expression profiles of effector CD8 T cells under normoxic conditions, where an active, effector signature is prominent, contrast with the hypoxic condition, showing reduced expression of these effector genes. Unpaired two-samples Wilcoxon test of significances are shown.

### Hypoxic T Cell Signature and Its Clinical Implications in Melanoma

Based on these observations, we sought to examine the hypoxia-induced T cell signature in melanoma clinical outcomes. Utilizing the CRI iATLAS melanoma RNAseq dataset,^31–36^ we examined the correlation of the hypoxia signature with clinical responses in samples from melanoma patients prior to immunotherapy treatment. We found that a hypoxic T cell signature was significantly correlated with reduced overall survival (p = 0.032) (Figure 5A) and progression-free survival (p = 0.0069) (Figure 5B). Separately, we extended our analysis to the ORIEN AVATAR cohort of melanoma patients undergoing ICI. We found that in gene expression data from patient samples prior to ipilimumab (IPI) and nivolumab (NIVO) treatment, the hypoxic T cell signature was associated with decreased survival rates, p = 0.042 and 0.041, respectively (Figures 5C and 5D). Altogether, the hypoxia T cell signature was predictive and consistently associated with patients with the worst clinical outcome. The consistent patterns across different datasets and treatment modalities underscore the clinical significance of hypoxia in T cells prior to ICI treatment therapy.

**Figure 5:**
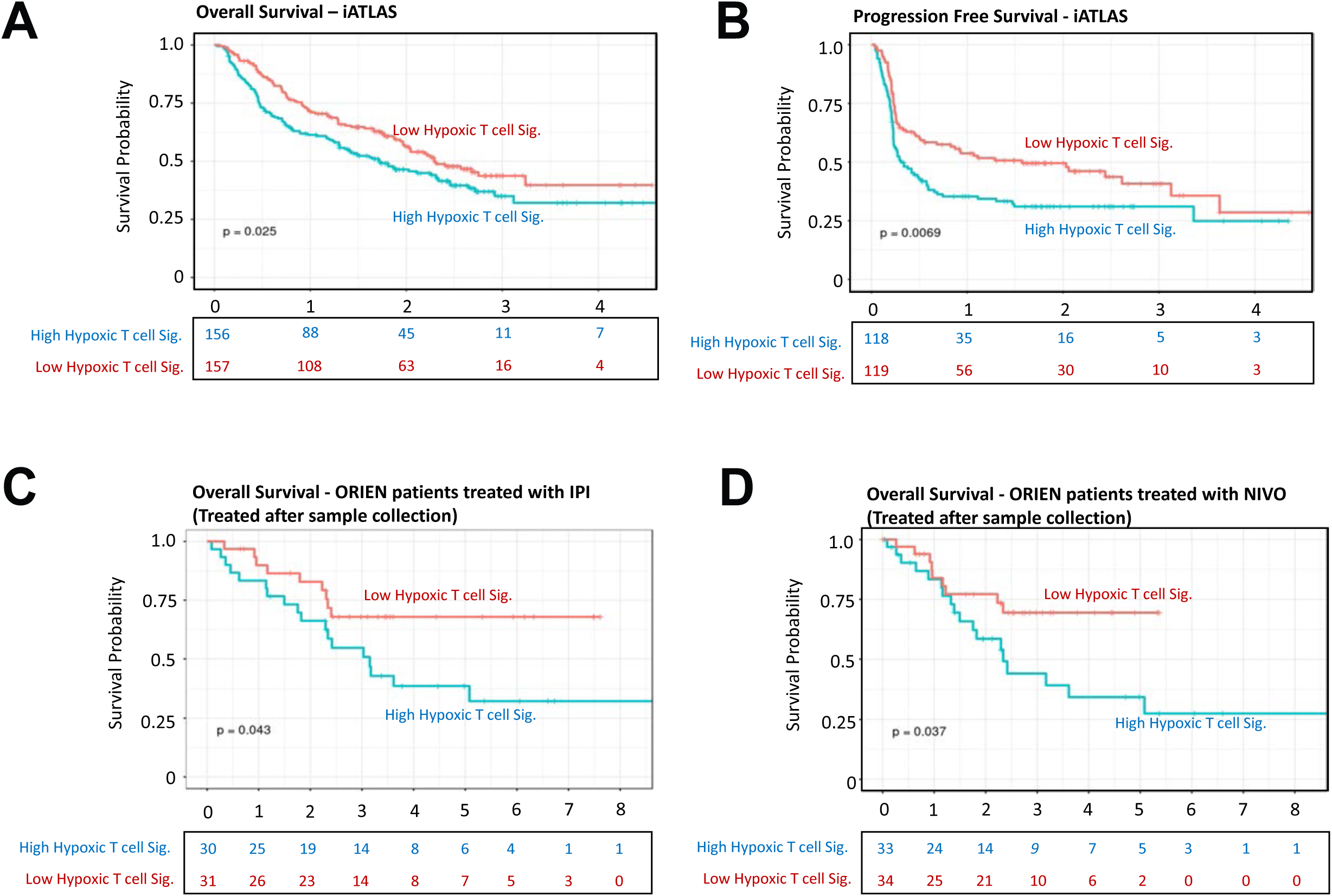
Hypoxic T Cell Signature and Its Clinical Implications in Melanoma. **(A)** Overall survival analysis of melanoma patients from the iATLAS dataset. A high hypoxic T cell signature correlates with significantly reduced overall survival. **(B)** Progression-free survival analysis showing lower rates in patients with a high hypoxic T cell signature. **(C-D)** Survival analysis of melanoma patients from the Moffitt ORIEN cohort treated with Immune Checkpoint Inhibitors (ICI). A high hypoxic T cell signature correlates with decreased survival rates in patients treated with Ipilimumab (IPI) and Nivolumab (NIVO). Survival was estimated based on the Kaplan-Meier method.

### T Cell Hypoxia-Signature Resembles “CD4/CD8 (T_STR_ States)”

Our initial analysis compared T cell signatures with total tumor RNA across multiple datasets to understand tumor-T cell interactions. Building on this, we conducted a more detailed examination by aligning our hypoxia signature with single-cell RNA sequencing (scRNA) data of T cells from tumors, utilizing the TCellMap database assembled by L. Wang’s group^16^ as a reference. Using the UMAP technique, we examined the distribution of CD8 and CD4 T cells categorized by their stress levels (Stressed vs. Not Stressed, (T_STR_ States)) and oxygenation status (Hypoxic vs. Normoxic) within the tumor microenvironment (Figure 6A, C).

**Figure 6:**
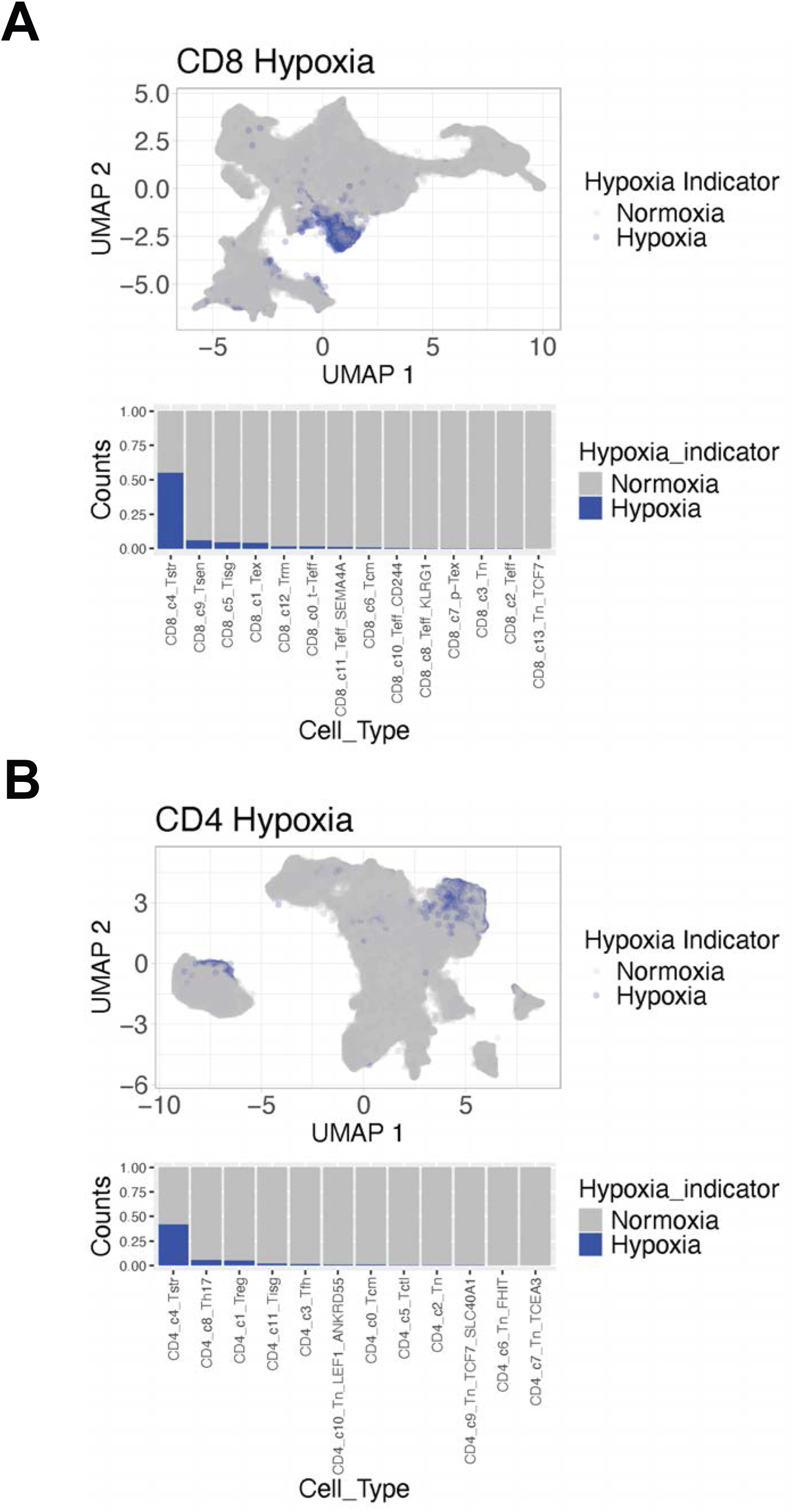
T Cell Hypoxia-Signature Resembles “CD4/CD8 Stressed Tumor T cells (TSTR States)” **(A, B)** Under hypoxic conditions, 2943 CD8 T cells were stressed versus 1923 not stressed. Under normoxic conditions, 2376 CD8 T cells were stressed versus 102,976 not stressed. Pearson’s Chi-squared test (χ² = 34,319, df = 1) and Fisher’s Exact Test (p < 2.2e-16, OR = 66.4, 95% CI: 61.7 to 71.1) indicate a significant association, with higher odds of CD8 T cells being stressed under hypoxia. **(C, D)** Under hypoxic conditions, 6271 CD4 T cells were stressed versus 3183 not stressed. Under normoxic conditions, 8666 CD4 T cells were stressed versus 153,641 not stressed. Pearson’s Chi-squared test (χ² = 41,848, df = 1) and Fisher’s Exact Test (p < 2.2e-16, OR = 34.9, 95% CI: 33.3 to 36.7) indicate a significant association, with higher odds of CD4 T cells being stressed under hypoxia.

Under Hypoxic conditions, 2943 CD8 T cells were categorized as Stressed, while 1923 were Not Stressed. In contrast, under Normoxic conditions, 2376 CD8 T cells were Stressed, with a substantial 102976 categorized as Not Stressed. Statistical analyses showed that Pearson’s Chi-squared test with Yates’ continuity correction had a chi-squared statistic of 34319 with 1 degree of freedom, indicating a highly significant association (p-value < 2.2e-16). Fisher’s Exact Test confirmed the association, with a p-value < 2.2e-16 and an odds ratio of 66.39914 (95% CI: 61.72125 to 71.10298), suggesting significantly higher odds of CD8 T cells being stressed under Hypoxic conditions compared to Normoxic conditions, (Figure 6A, B).

Similarly, under Hypoxic conditions, 6271 CD4 T cells were categorized as Stressed, while 3183 were Not Stressed. In contrast, under Normoxic conditions, 8666 CD4 T cells were Stressed, with a substantial 153641 categorized as Not Stressed. Statistical analyses showed that Pearson’s Chi-squared test with Yates’ continuity correction had a chi-squared statistic of 41848 with 1 degree of freedom, indicating a highly significant association (p-value < 2.2e-16). Fisher’s Exact Test confirmed the association, with a p-value < 2.2e-16 and an odds ratio of 34.91855 (95% CI: 33.28564 to 36.68525), suggesting significantly higher odds of CD4 T cells being stressed under Hypoxic conditions compared to Normoxic conditions, (Figure 6C, D).

These findings underscore the pronounced influence of oxygenation status on CD4 and CD8 T cell stress levels within the TME, highlighting potential implications for understanding immune responses in cancer contexts.

## DISCUSSION

This study demonstrates the critical influence of hypoxia on T cell distribution within the TME, highlighting its profound impact on T cell function and status, and its clinical implications. Using oxygenation and T cell accessibility maps derived from melanoma TMAs, we reveal a detailed spatial relationship between T cells and oxygen availability. Importantly, we identify a distinct hypoxic gene signature in T cells under low oxygen conditions, which correlates with stressed and inactive T cells and poorer prognosis in melanoma patients, particularly among those who do not respond to ICIs.

Our observations of T cell spatial distribution and oxygenation levels in melanoma TMAs indicate that T cells tend to accumulate close to vasculature in highly oxygenated areas, suggesting that oxygen availability is a crucial factor in their localization and anti-tumor function. These findings have significant implications for understanding how T cells exert their cytotoxic effects within the tumor. In cases where immunotherapy is effective, T cells might be able to penetrate the tumor from the periphery and exert their cytotoxic functions. However, it remains an open question whether T cells primarily kill tumor cells from the inside or the outside of the tumor mass. Interestingly, while in cores the number of T cells is in thousands and in peripheries they number in hundreds, we observed no differences in T cell distribution between core and peripheral biopsies, suggesting a uniform influence of oxygenation across these regions.

We found that in some cases, CD4 T cells locate in hypoxic areas. While our staining strategy did not include FoxP3, it is possible that these are regulatory T cells (Tregs). Studies indicate that CD4 T cells, including Tregs, exhibit specific distribution patterns within the TME that are significantly influenced by hypoxic conditions. Hypoxia plays a crucial role in promoting the differentiation and function of Tregs. Hypoxia-inducible factor 1-alpha (HIF-1α) can directly enhance the expression of FoxP3, a key transcription factor for Treg development, thereby increasing the number of Tregs in hypoxic regions.^37^

An important study by Gallon et al.^10^ examined the relationship between the type, density, and location of TILs and clinical outcomes in colorectal cancer patients. They observed that higher densities of CD3, CD8, GZMB, and CD45RO cells correlate with improved PFS and OS. Importantly, their study did not specifically investigate how hypoxia correlates with these cell types and their distribution within the TME, focusing instead on the general presence and characteristics of immune cells in the TME. Our study also underscores the critical role of immune cells in influencing clinical outcomes. Future research should aim to elucidate the precise mechanisms and spatial dynamics of T cell-mediated tumor cell killing, particularly concerning varying oxygen levels within the TME. This understanding could inform strategies to enhance T cell infiltration and function in hypoxic tumor regions, potentially enhancing the efficacy of immunotherapies.

At the transcriptional level, hypoxic T cells displayed significant alterations in gene expression. Our RNA sequencing analysis revealed that hypoxic T cells adopt a gene expression profile similar to that of resting or undifferentiated T cells, contrasting with the active effector T cell profile observed under normoxic conditions. Specifically, genes such as FOS and FOSB, crucial for immune cell development and cytokine production, were significantly downregulated under hypoxia.^38, 39^ IFN-γ, critical for T cell activation and promoting anti-tumor immunity, was also significantly reduced.^40, 41^ Additionally, VEGFA was upregulated under low oxygen conditions, promoting angiogenesis and reflecting adaptive responses to hypoxic stress.^3^ Other genes impacted by hypoxia include CCL3 (MIP-1α), FADS1, GRM2, IL17F, INSIG1, LDLR, and SCD, which play various roles in immune cell recruitment, lipid metabolism, and inflammation.

Our findings align with several previous studies that have explored the impact of hypoxia on immune cells and the TME. Wigerup et al.^11^ identified a seven-gene hypoxia signature associated with worse outcomes in cutaneous melanoma, supporting our observations on the negative prognostic impact of hypoxia. Buffa et al. and Eustace et al.^42, 43^ developed hypoxia metagenes predictive of poor prognosis and therapy resistance in various cancers, similar to our findings in melanoma. Ye et al.^44, 45^ demonstrated that hypoxia-induced transcriptional changes in lung cancer led to immune suppression and poor clinical outcomes, paralleling our observations. Toustrup et al.^46^ identified a hypoxia gene expression classifier predicting survival in head and neck cancer, reinforcing the value of hypoxia signatures as prognostic markers across cancers.

However, these studies consistently focus on whole tumors that include tumor cells and immune components. In contrast, our study specifically emphasizes the hypoxic effects on T cells. By isolating and examining the T cell population, we highlight the unique transcriptional and functional impairments induced by hypoxia, which are distinct from the broader tumor hypoxia effects observed in previous research. This targeted approach allows us to pinpoint how hypoxia directly influences T cell efficacy within the TME, providing a more detailed understanding of immune evasion mechanisms at the cellular level.

The clinical implications of these findings are profound, suggesting that hypoxic conditions within the tumor microenvironment significantly impact T cell function and distribution, leading to poorer clinical outcomes in melanoma patients treated with ICI. Specifically, our analysis using the iATLASdataset revealed that a hypoxic T cell signature is associated with reduced overall survival and progression-free survival. This was further validated in the Moffitt ORIEN cohort, where patients with this hypoxic T cell signature showed decreased overall survival rates during ICI therapy. These consistent patterns across various datasets and treatment modalities highlight the prognostic value of the hypoxic T cell signature. The implications include using the hypoxic T cell signature as a prognostic marker, targeting hypoxic regions within tumors or modulating hypoxia-induced pathways to enhance immunotherapy, developing personalized treatment approaches for patients with a high hypoxic signature, and directing future research towards therapeutic interventions that address hypoxia-induced T cell dysfunction. Overall, considering hypoxia in the tumor microenvironment when developing cancer immunotherapies could significantly improve patient outcomes in melanoma and other cancers.

Chu et al.^16^ have identified a unique T cell stress response state (T_STR_), characterized by heat shock gene expression, within the tumor microenvironment across various cancer types. This T_STR_ state is associated with immunotherapy resistance, particularly in non-responsive tumors. Similarly, our study demonstrates that hypoxia significantly impacts T cell distribution, function, and gene expression, resulting in a hypoxic T cell signature associated with poorer clinical outcomes in melanoma patients. Using the UMAP technique, we examined the distribution of CD8 and CD4 T cells categorized by their stress levels (Stressed vs. Not Stressed) and oxygenation status (Hypoxic vs. Normoxic) within the tumor microenvironment, revealing significant findings through statistical analyses.

Our findings show that under hypoxic conditions, there is a significantly higher prevalence of stressed CD8 and CD4 T cells compared to normoxic conditions. Specifically, Pearson’s Chi-squared test with Yates’ continuity correction and Fisher’s Exact Test revealed highly significant associations, with the odds of T cell stress being markedly higher under hypoxia. These findings underscore the pronounced influence of oxygenation status on T cell stress levels within the tumor microenvironment, highlighting potential implications for understanding immune responses in cancer contexts.

The pronounced stress response observed in T cells under hypoxia can be linked to their impaired functionality, as evidenced by altered gene expression profiles. Genes such as FOS, FOSB, and IFN-γ, which are critical for T cell activation and cytokine production, were significantly downregulated, aligning with the hypoxic stress response observed in our study. This impaired function likely contributes to the reduced efficacy of T cells in targeting and killing tumor cells in hypoxic regions, thereby affecting overall clinical outcomes.

While reducing oxygen to the tumor site might initially seem beneficial for starving the tumor, it also risks impairing the immune response. Enhancing or normalizing tumor vasculature to improve oxygenation could be a promising strategy to boost immune responses while simultaneously making it harder for cancer cells to adapt and survive. This approach might offer a more balanced and effective way to target tumors, especially when integrated with immunotherapies. Several other studies also support the prognostic and therapeutic implications of targeting hypoxia in cancer treatment. For instance, the study by Vaupel et al.^47^ highlights that tumor hypoxia is associated with resistance to conventional therapies and a poorer prognosis. This is in line with our findings that hypoxia contributes to T cell dysfunction and immune evasion. Moreover, research by Minchenko et al. ^48^ discusses the metabolic adaptations of tumor cells under hypoxia and the potential for increased oxidative stress upon reoxygenation, suggesting that reoxygenation following hypoxia can increase the vulnerability of tumor cells to oxygen-mediated cell death.

Future research should focus on developing therapeutic strategies targeting hypoxia-induced pathways in T cells. Understanding the mechanisms behind hypoxic stress responses in T cells is crucial for effective interventions. Investigating how hypoxia drives T cells into a less active state and identifying targets to reverse this effect could lead to novel immunotherapies. Engineering T cells to resist hypoxic stress or enhancing their metabolic adaptability could improve efficacy in hypoxic TMEs. Additionally, integrating transcriptional profiling with clinical data can identify biomarkers for immunotherapy response, helping tailor approaches to overcome resistance and improve outcomes.

In conclusion, our study reveals critical insights into the profound influence of hypoxia on T cell distribution, function, and gene expression within the TME of melanoma. We demonstrate that hypoxic conditions lead to a distinct T cell gene signature associated with compromised anti-tumor efficacy and poorer clinical outcomes in melanoma patients, including those receiving ICI therapy. These findings underscore the need to consider hypoxia as a pivotal factor in designing effective cancer immunotherapies. By targeting hypoxia-induced pathways in T cells, such as enhancing metabolic adaptability or engineering resilience to hypoxic stress, novel therapeutic strategies could be developed to improve treatment responses and patient outcomes in melanoma and other cancers. This comprehensive understanding of hypoxia-mediated immune evasion mechanisms highlights promising avenues for future research and clinical translation.

---

**Figure S1 Legend**

**Spatial Distribution and Oxygenation of CD4 and CD8 T Cells in Human Melanoma Tissue Microarrays (TMAs). (A-B)**: Fractions of CD8 and CD4 T cells for 17 core regions of the melanoma TMAs-T cells were quantified for their distance from the nearest blood vessel **(A)**, and oxygen level in their vicinity **(B)**. **(C-D)**: Fractions of CD8 and CD4 T cells for 13 peripheral regions of the melanoma TMAs-similarly, T cells were analyzed for their distance from blood vessels **(C)**, and oxygen level in their vicinity **(D)**. In all cases, the majority of T cells are localized within 50-100 micrometers from the vasculature in the regions of high oxygen availability.

## Declarations

### Ethics Approval and Consent to Participate

The study was conducted under the approval of the Institutional Review Board (IRB) at Moffitt Cancer Center, ensuring adherence to ethical guidelines and patient consent protocols.

## Consent for Publication

### Availability of Data and Material

The data that support the findings of this study are available from the corresponding author upon reasonable request.

### Funding

This research was funded the NIH/NCI grants R01CA250514, and the VoLo Health Foundation (to JAG) and by R01CA259387 and R01CA272601 (to KAR). The study was funded by the ICADS T32 postdoctoral training program at the Moffitt Cancer Center (NCI T32 CA233399) (to JD) and P30-CA076292 to the Moffitt core facilities. This work has been supported in part by the Biostatistics and Bioinformatics Shared Resource at the H. Lee Moffitt Cancer Center & Re-search Institute, an NCI-designated Comprehensive Cancer Center.

### Authors’ Contributions

Mate Z. Nagy: Conceptualization, Methodology, Writing - Original Draft

Lourdes B. Plaza Rojas: Conceptualization, Methodology, and Analysis

Justin C. Boucher: Writing - Review & Editing

Elena Kostenko: Validation, Methodology

Anna L. Austin: Validation, Methodology

Joshua Davis: Validation, Methodology

Zhihua Chen: Validation, Methodology

Dongliang Du: Validation, Methodology

Awino Maureiq E. Ojwang: Validation, Methodology

Ahmad Tarhini: Clinical input and ORIEN

Katarzyna A. Rejniak: Methodology, Mathematical Modeling and Analysis

Timothy Shaw: Methodology, Bioinformatics Analysis

Jose Alejandro Guevara: Conceptualization, Funding Acquisition, Writing - Review & Editing

## Acknowledgements

We thank the staff at Moffitt Cancer Center for their support and assistance with this study. Special thanks to the Cancer Center’s Analytical Microscopy Core and Joseph Johnson for his assistance, and to Carlos M. Morán-Segura from the Histology Core, the Cancer Center’s Flow Cytometry Core and Jodi Kroeger as well as Alyssa Obermayer for their contributions.

## List of Abbreviations

CAR: Chimeric Antigen Receptor
CTV: Cell Trace Violet
GSEA: Gene Set Enrichment Analysis
ICI: Immune Checkpoint Inhibitors
IFN-γ: Interferon-Gamma
IRB: Institutional Review Board
IPI: Ipilimumab
NIVO: Nivolumab
ORIEN: Oncology Research Information Exchange Network
OS: Overall Survival
RNAseq: RNA Sequencing
scRNA: Single-Cell RNA Sequencing
TCGA: The Cancer Genome Atlas
TILs: Tumor-Infiltrating Lymphocytes
TME: Tumor Microenvironment
UMAP: Uniform Manifold Approximation and Projection

