## Supplementary figures and images for "Effector T Cells under Hypoxia have an Altered Transcriptome Similar to Tumor-Stressed T Cells Found in Non-Responsive Melanoma Patients"

### Figure S1

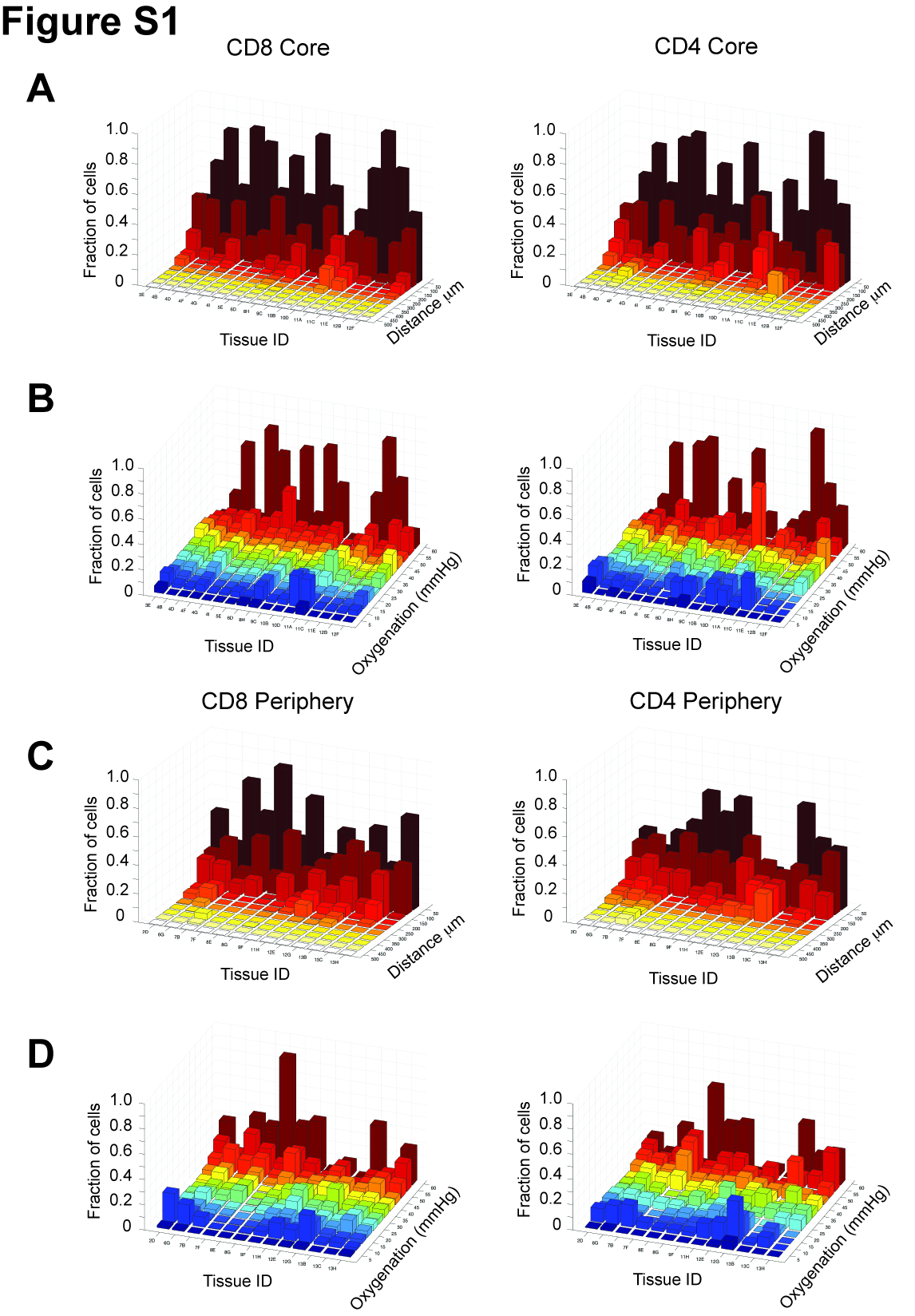
